# Epithelial folding through local degradation of an elastic basement membrane plate

**DOI:** 10.1101/2024.02.06.579060

**Authors:** K. Yanín Guerra Santillán, Caroline Jantzen, Christian Dahmann, Elisabeth Fischer-Friedrich

**Affiliations:** Cluster of Excellence Physics of Life, Technische Universität Dresden, Dresden, Germany; School of Science, Technische Universität Dresden, Dresden, Germany; Biotechnology Center, Technische Universität Dresden, Dresden, Germany

## Abstract

Epithelia are polarised layers of cells that line the outer and inner surfaces of organs. At the basal side, the epithelial cell layer is supported by a basement membrane, which is a thin polymeric layer of self-assembled extracellular matrix (ECM) that tightly adheres to the basal cell surface. Proper shaping of epithelial layers is an important prerequisite for the development of healthy organs during the morphogenesis of an organism. Experimental evidence indicates that local degradation of the basement membrane drives epithelial folding. Here, we present a coarse-grained plate theory model of the basement membrane that assumes force balance between i) cell-transduced active forces and ii) deformation-induced elastic forces. We verify key assumptions of this model through experiments in the *Drosophila* wing disc epithelium and demonstrate that the model can explain the emergence of outward epithelial folds upon local plate degradation. Our model accounts for local degradation of the basement membrane as a mechanism for the generation of epithelial folds in the absence of epithelial growth.

## I. INTRODUCTION

During the morphogenesis of an embryo, precursor tissues need to undergo complex shape changes to develop into their final functional form in an adult organism^1–4^. Epithelia are sheet-like arrays of cells that line the surface of organs or body cavities. They are key components of organ function^2,5^. A major element of epithelial morphogenesis is epithelial folding^2,5^, which is particularly prominent in the development of the epithelium of our brain or intestine^3^. A prevalent biological model organism used to study epithelial morphogenesis is the fruit fly *Drosophila melanogaster*. There, epithelial folding is part of the healthy morphogenesis of the columnar epithelium of the larval wing imaginal disc, the precursor tissue of the adult fruit fly wing.

Epithelial cell sheets are polarized with an apical and basal surface. The basal epithelial surface is attached to a thin layer of specialized extracellular matrix called basement membrane with a typical thickness between tens of nanometers to several micrometers^6,7^. Locally emerging differences in apical and basal tension of the epithelial cell faces were identified as major driving mechanism of epithelial folding. While apical tension is mainly regulated through active molecular myosin motors in the actin cytoskeleton attached to the apical cell faces, we recently identified passive elastic stresses from elastic basement membrane stretch as a major contributor to basal tension^8^.

Cellular mechanics in mammalian epithelia was shown to be dominated by the intracellular actin cytoskeletal network for fast time scales^9,10^. However, for larger time scales, actin-cytoskeleton-related cell stiffness decreases with an increasingly fluid-like nature of the material^11–13^. Correspondingly, deformation-induced elastic stresses in the actin cytoskeleton dissipate on longer time scales. It has been suggested that this mechanical trend is due to fast turnover of molecular components in the actin cytoskeleton^10,14–16^. By contrast, the basement membrane at the basal face of the epithelium was shown to bear elastic stresses also on longer time scales of several minutes and beyond^8,17–19^. Furthermore, the basement membrane was shown to behave solid-like on time scales of hours, i.e. it can be considered as a solid-like material in the course of morphogenetic changes such as epithelial folding^8,16,20^. It must therefore not astonish that basement membranes play a major role in the mechanical stabilization and the shape regulation of tissues^6,17,19,21^. In particular, previous work has reported that local degradation of the basement membrane can induce epithelial folding in the columnar epithelium of the wing disc in the 3rd instar larva of the fruit fly *Drosophila melanogaster*^21^. We note, that in this same study a 3D vertex model was employed to account for apical folding upon local basement membrane degradation. However, assuming a permanently flat basement membrane, basal folding was explicitly not incorporated in this modelling approach^21^.

Here, we propose a continuum-mechanical model to rationalize the observation of basal epithelial folding upon local degradation of the basement membrane. While already previous studies have used plate theory to account for folding upon local plate degradation in epithelia^22,23^, these models were relying on epithelial growth as a key mechanism for folding. However, it was shown experimentally that formation of the H/H fold in the fruit fly larval tissue takes place in a timely manner in the absence of cell division^21^ suggesting that fold formation is independent of tissue growth.

Correspondingly, our model strives to account for basal folding relying on a growth-independent mechanism; for this purpose, we focus on the emergent shape of the solid-like basement membrane which we model as an elastic plate in contact with an epithelial cell layer that exerts external forces on the basal plate. Our model anticipates an elastic prestress balanced by an active in-plane stress exerted by the neighboring cell layer. The combination of opposing active and passive prestress contributions allows for an initially flat plate conformation which turns into a folded conformation upon perturbation of the existing force balance by local basement membrane degradation. Upon folding onset, an orthogonal net force emanating from hydrostatic pressure excess in the epithelial cells emerges in the fold region and further increases fold depth. In this way, we present a model that describes the initiation of basal epithelial folding towards the basal side. The associated force balance equations are solved analytically in the framework of linear elasticity theory. Key predictions of the model such as the role of hydrostatic pressure and of basal tension in regulation of fold depth are verified experimentally in folds of the wing disc epithelium.

## II. THEORETICAL MODEL OF WING DISC FOLD FORMATION

Our plate model focuses on the wing disc mid region (region in the center next to the A-P boundary^24^, see black box in Fig. 1a). In this region, the basement membranes of the columnar epithelium and of the peripodial epithelium are modeled by segments of thin plates (blue and green) connected by linker elements (red) with semi-circular cross-sections (Fig. 1b,e). For simplicity, we anticipate that the linker elements are rigid in nature and do not deform during fold formation. Furthermore, we consider only the central stripe along the wing disc long axis (region in the center next to the A-P boundary indicated by the black box in Fig. 1a). In addition, we assume that there is no curvature and no change of wing disc mechanical parameters along the y-axis.

**Figure 1.**
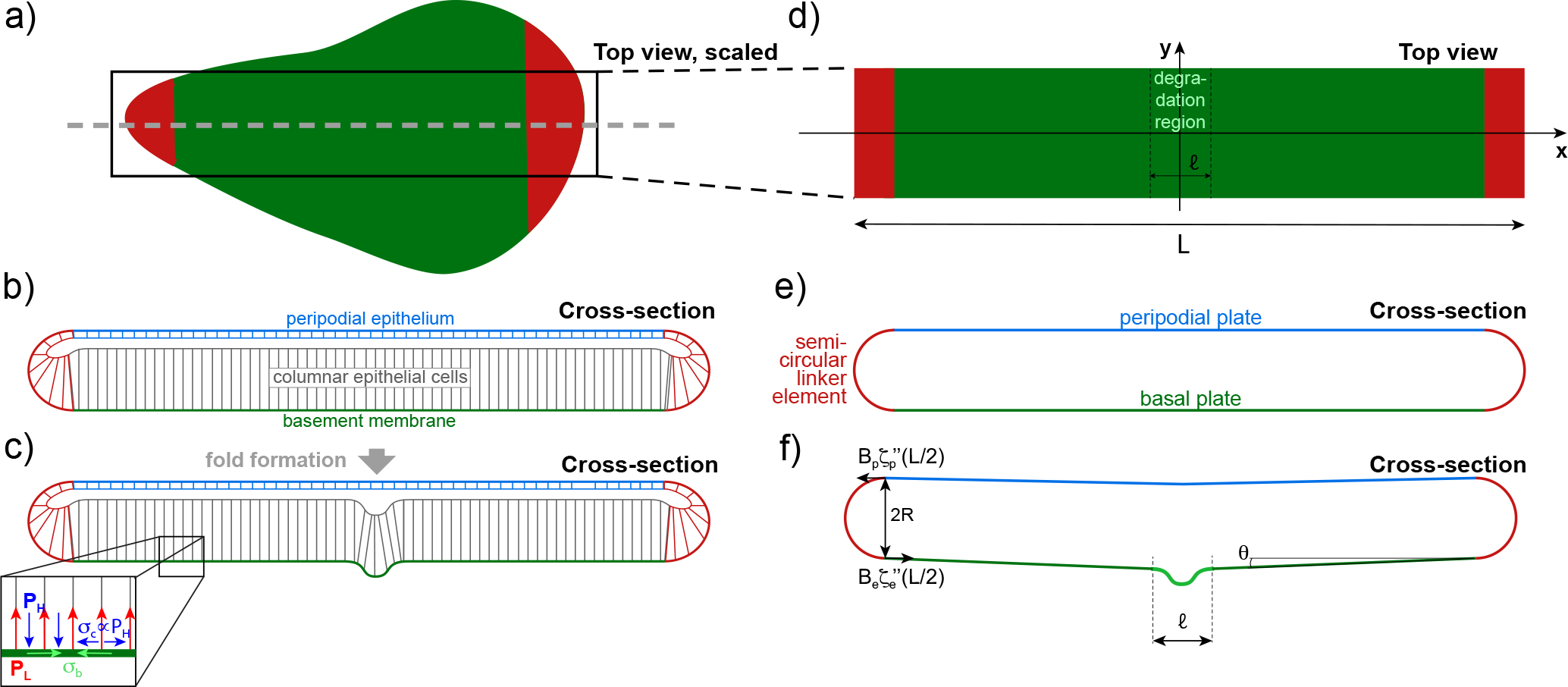
Schematic of wing disc architecture before and after folding (a-c) and corresponding mechanical elements of the plate model (d-f). a) Scaled, basal view of the wing disc. The grey dashed line indicates where the cross section in panel b is taken. The black box outlines the region taken into consideration for theoretical modelling as shown in panel e. b) Lateral view of wing disc cross-section before folding. c) Lateral view of the wing disc cross-section after the first fold (H/H fold) has formed. d) Basal view of the rectangular plate model. e) Lateral view of the plate model before folding. The blue and green plate segments model the basement membrane in the peripodial epithelium and in the columnar epithelium, respectively. The red semi-circular pieces are rigid mechanical linker elements. f) Lateral view of the plate model after folding. The angle *θ* quantifies the angle between the plate tangent and the horizontal axis at the boundary between green and red plates.

In the unperturbed state, we anticipate that our plate’s extension is in the x-y plane (Fig. 1a,b,d,e). Furthermore, we assume that i) external tangential net forces are zero, ii) stress, strain and deformation components are constant along the y-axis (apart from a possible linear change of the y-component of the in-plane displacement vector *u*_*y*_), and iii) curvature components *C*^*xy*^ and *C*^*yy*^ are zero. Let the function *ζ*(*x*) denote the displacement of the neutral line of the plate in z-direction. Note that the curvature of the plate *C*^*xx*^ is to first order given by −*ζ*^*′′*^(*x*). Correspondingly, the bending moment at position *x* is *m*_*xy*_ = −*Bζ*^*′′*^(*x*), where *B* is the plate’s bending modulus.

Coarse-graining over the scale of cell widths, we anticipate that there are two isotropic in-plane prestress contributions in the basal plate: i) a contractile prestress *σ*_*b*_(*t*) resulting form elastic basement membrane stretching and ii) an active expansile prestress generated by the hydrostatic pressure excess in the neighbouring cell layer *σ*_*c*_(*t*) (Fig. 1c). Before fold formation (*t* = 0), we assume that there is a balance of active and passive prestress such that

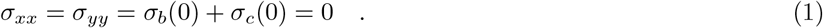

We divide the basal layer of the model wing disc into a degradation region extending from *x* = −*𝓁/*2 to *x* = *𝓁/*2 and a surrounding external region spanning the intervals [ −*L/*2, − *𝓁/*2] and [*𝓁/*2, *L/*2] (Fig. 1e,f). We will anticipate that due to basement membrane degradation *σ*_*b*_ is reduced over time in the internal degradation region by a constant stress amount Δ*σ*(*t*), i.e. *σ*_*b*_(*t*) = *σ*_*b*_(0) −Δ*σ*(*t*). We assume that this change in prestress is slow as compared to the equilibration of the viscoelastic dynamics. Therefore, we presuppose a quasi-static state of the wing disc at all times while Δ*σ*(*t*) is slowly increasing. Accordingly, we will omit the argument *t* in Δ*σ*(*t*) in the forthcoming calculations. Further, we anticipate that the expansile prestress in the cell layer does not change over time. Using these assumptions, we will, in the following, formulate the governing continuum-mechanical equations of plate deformation for small strain. To keep the force balance during basal prestress degradation, the layers of the wing disc deform elastically. Since the amount of contractile prestress in the collagen is reduced, there will be an expansile deformation in the degradation region in response.

After deformation, the out-of plane deflection *ζ*(*x*) of the plate contributes an in-plane stretch^25^ *ζ*^*′*^(*x*)^2^*/*2. Further, there can be an additional in-plane stretch quantified by a horizontal displacement vector^25^ (*u*_*x*_, *u*_*y*_). Correspondingly, the xx-component of the strain reads *u*_*xx*_ = *∂*_*x*_*u*_*x*_(*x*) + *ζ*^*′*^(*x*)^2^*/*2. With this notation, the xx- and yy-component of the resulting stress tensor in the different regions of our model wing disc read^25^

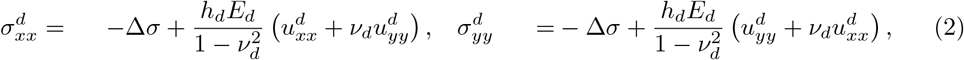

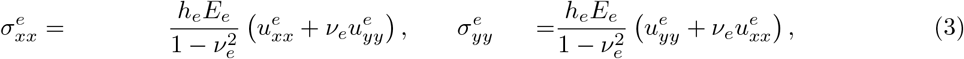

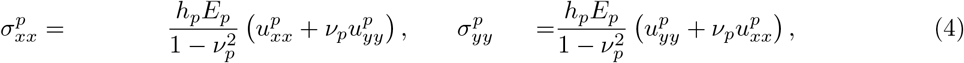

where the sub/superscripts *d, e* or *p* mark variables adopted in the basal degradation region, the basal external region or in the peripodial plate, respectively. The elastic parameters *ν* and *E* are defined as *ν* = (*K*_*B*_ −*K*_*S*_)*/*(*K*_*B*_ + *K*_*S*_) and *E* = 2*K*_*B*_(1 −*ν*)*/h*, where *K*_*B*_, *K*_*S*_ and *h* are the area bulk modulus, the area shear modulus and the thickness of the plate, respectively. (In the case of a thin plate made of an isotropic material, *ν* is the Poisson ratio and *E* is the Young’s modulus of the plate material^26^.) Note that 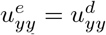, see Appendix A.

The tangential and normal force balance conditions require for every region of the model wing disc (see Appendix A)

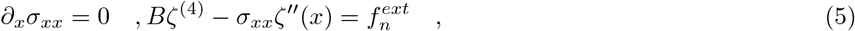

where the respective sub/superscript *d, e* or *p* has to be added to each variable in the respective region. The second equation also follows from the Föppl-von Kármán equations for the equilibrium of plates with large deflection^25,27^. Here, 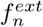 is the external normal force per unit area which is exerted by the neighbouring epithelial cells on the basement membrane plate. We assume that it has two contributions: i) a force density *P*_*L*_ exerted by contractility of the lateral cell walls which pulls the basement membrane towards the cell layer and ii) a force density −*P*_*H*_ generated by the hydrostatic pressure excess *P*_*H*_ that pushes the basement membrane away from the cell layer, see Fig. 1c, inset. Applying coarse-graining over the cell-scale, we anticipate, that there is a force balance such that 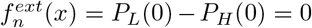 before fold formation. During fold formation, there is an in-plane stretch that leads to a change in density of lateral cell walls along the wing disc. Correspondingly, during fold formation, *P*_*L*_ is now to first order *P*_*L*_ = *P*_*L*_(0)(1 −*u*_*xx*_ −*u*_*yy*_). By contrast, we anticipate that the contribution *P*_*H*_ remains constant over time (Fig. 1a). Hence, we find 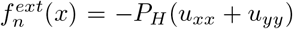 during fold formation. (It is noteworthy that *u*_*yy*_ is constant, see Appendix A, and therefore, also *u*_*xx*_ is constant within one region in our model for a step-wise increase of Δ*σ* in the degradation region.)

In the following, we will make the simplifying assumption that the degradation region is narrow such that the remaining plate regions dominate mechanically. In this scenario, stretch energy minimization will lead to *σ*_*xx*_ ≈ *σ*_*yy*_ ≈ 0 at all time points outside the degradation region. Correspondingly, *σ*_*xx*_ and *u*_*yy*_ are also approximately zero everywhere including the degradation region. With this, we obtain with Eq. (5) as final force balance condition for normal forces

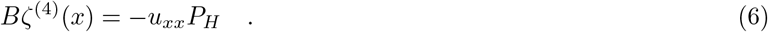

## III. SOLUTIONS TO PLATE SHAPE AND IN-PLANE DEFORMATION

Solving Eq. (6), we make the following functional ansatz for the function *ζ*(*x*) in the three different regions of the model wing disc using Eqn. (6) and (2)-(4) with *σ*_*xx*_ = 0

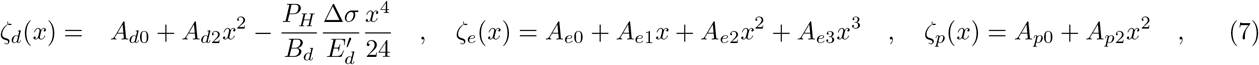

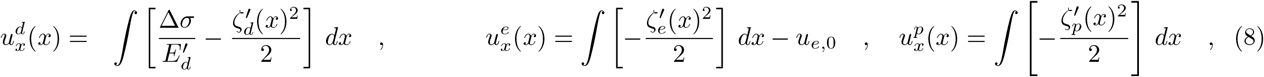

where antiderivatives in Formulae (8) are taken such that displacements 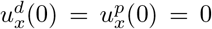. Furthermore, *E*^*′*^ = *hE/*(1 −*ν*^2^). One may check that this ansatz automatically fullfils the force balance conditions given by Eq. (6) in the two regions of the basal plate and in the peripodial plate. The constants *A*_*d*0_, *A*_*d*2_, *A*_*e*0_, *A*_*e*1_, *A*_*e*2_, *A*_*e*3_, *A*_*p*0_, *A*_*p*2_ and *u*_*e*,0_ need to be fixed by the boundary conditions given in rows 1-9 in Table I.

**Table I.** Boundary conditions for Eqn. (7) and (8).

| # | boundary condition | reason for the boundary condition |
| --- | --- | --- |
| 1 | $\zeta_p(L/2) = \zeta_e(L/2) + 2R\cos(\theta)$ | z-coordinate matching of basal and peripodial layer end points with the position of the end points of the rigid semi-circular linker elements |
| 2 | $\zeta'_p(L/2) = \zeta'_e(L/2)$ | matching of the tangents of the peripodial and basal plate layers at connection of the rigid semi-circular linker elements |
| 3 | $B_e\zeta''_e(L/2) + B_p\zeta''_p(L/2) + 2\sigma_{xx}R = 0$ | rigid semi-circular linker elements are torque-free |
| 4 | $\zeta_e(\pm l/2) = \zeta_d(\pm l/2)$ | z-coordinate matching of the plate layers at the interface of the internal degradation region and the external basal layer |
| 5 | $\zeta'_e(\pm l/2) = \zeta'_d(\pm l/2)$ | matching of the tangents at the interface of the internal degradation region and the external basal layer |
| 6 | $B_e\zeta''_e(\pm l/2) = B_d\zeta''_d(\pm l/2)$ | torque equilibrium at the interface of the internal degradation region and the external basal layer |
| 7 | $\zeta_e(\pm L/2) = 0$ | no z-displacement at $x = L/2$ |
| 8 | $u_p(L/2) = u_e(L/2) - 2R\sin(\theta)$ | x-coordinate matching of basal and peripodial layer end points with the position of the end points of the rigid semi-circular linker elements |
| 9 | $u_d(l/2) = u_e(l/2)$ | matching in-plane displacements at the interface of the internal degradation region and the external basal layer |
| 10 | $u_d(0) = 0, \quad u_p(0) = 0$ | no in-plane displacement at the origin |
| 11 | $\sin(\theta) = \zeta'_e(L/2), \cos(\theta) = 1 - \zeta'_e(L/2)^2/2$ | angle estimate up to first order |

To generate quantitative solutions for plate shapes of varying degrees of local basement membrane degradation, we need to set appropriate parameters for the model. Assuming that the basement membrane material is initially the same in all regions of the wing disc, plate elastic moduli are chosen such that the basement membrane material properties are equal everywhere before local degradation of the basement membrane starts. With increasing degradation degree *α*, the basement membrane prestress in the degradation region is assumed to cease such that elastic moduli in this region goes down. Correspondingly, we set 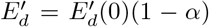 and *B*_*d*_ = *B*_*d*_(0)(1 − *α*)^3^, where *α* is the degree of plate degradation running from zero to one. (The cubic dependence of *B*_*d*_ on (1 −*α*) for the bending modulus is motivated by the cubic dependence of this modulus on plate thickness.) Here, *σ*_*b*_(0), *E*_*d*_(0) and *B*_*d*_(0) are the elastic prestress, the modulus *E*^*′*^ = *hE/*(1 −*ν*^2^) and the bending modulus in the degradation region before local basement membrane degradation starts. Parameter choices were motivated by experimental measurements^8^. The basal elastic prestress in the basement membrane was measured to be *σ*_*b*_(0) ≈ 0.3 mN/m. For small strains, the basement membrane area bulk modulus *K*_*B*_ was estimated to be in the range ≈ 0.5 −3 mN/m, see^8^. At a Poisson ratio of ≈ 0.5, we expect a similar value of *E*^*′*^. Taking into account that collagen networks have been reported to be subject to strain-stiffening^28,29^, we chose *E*^*′*^ slightly larger as 5 mN/m (same value in all wing disc regions before degradation). The experimentally most undetermined parameter is the bending modulus *B*. The bending modulus of an entire epithelial layer was previously determined by Fouchard *et al*. as ≈ 400 · 10^−15^Nm^30^. Here, we choose the bending modulus of the basement membrane plate to be approximately one order of magnitude lower as *B* = 50 × 10^−15^ Nm. The hydrostatic pressure in interphase cells was previously estimated to range between 10 − 100 Pa in non-adherent HeLa cells^31^. Therefore, we choose *P*_*H*_ = 50 Pa.

Calculating out-of-plane deflections for the parameter choices mentioned above, we see that softening of the degradation region in conjunction with diminished elastic prestress leads to a bulging out of the basement membrane in the degradation region, i.e. it leads to an induction of folding (Fig. 2a,d, e-g). Both, out-of-plane deflections and in-plane displacements contribute to elastic plate strain *u*_*xx*_, see Fig. 2b,c.

**Figure 2.**
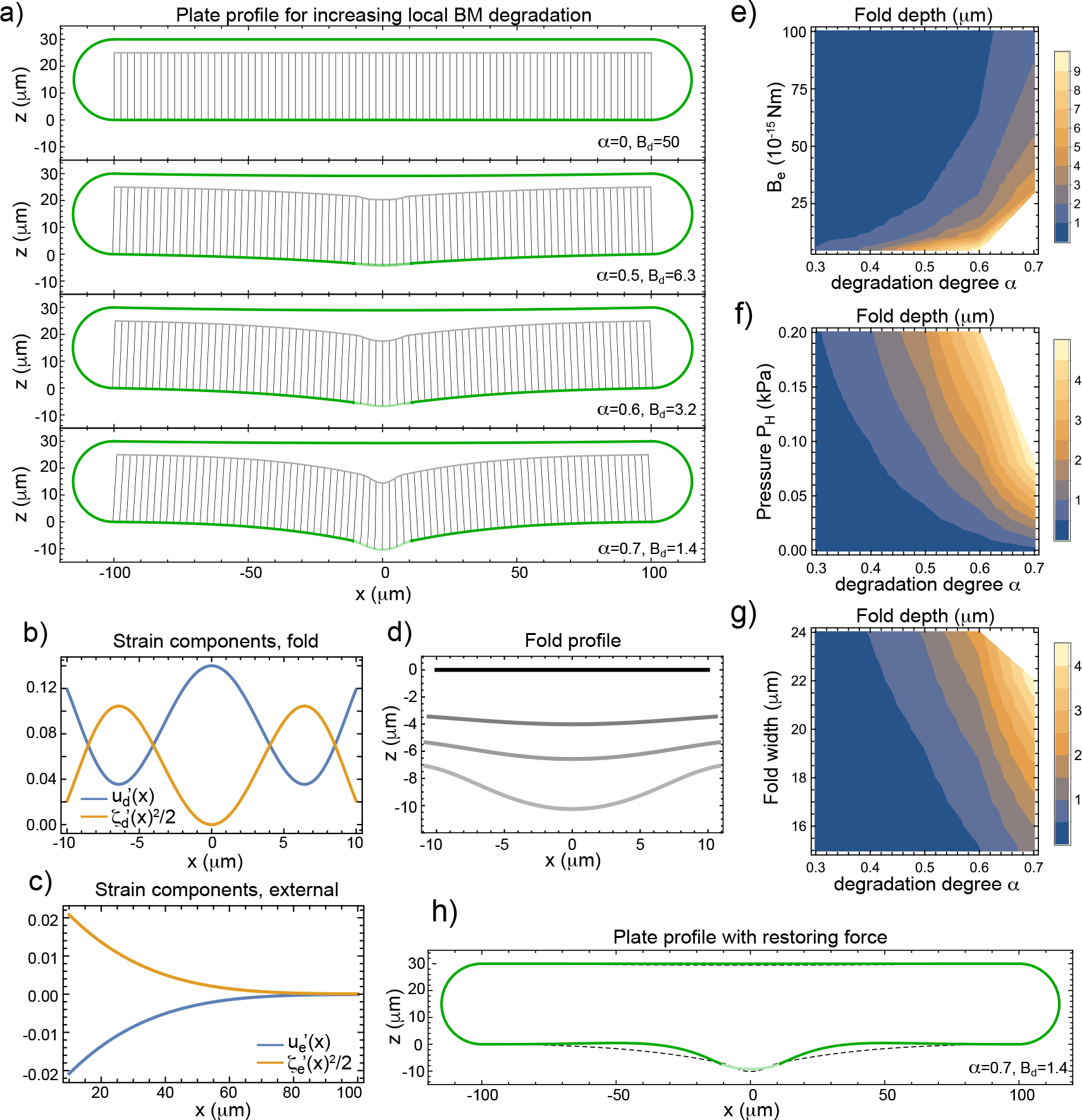
Wing disc folding in response to local basement membrane degradation. a) Plate shapes according to Eqn. (7) and (8) upon increasing local plate degradation in the central degradation region, see also Supplemental Movie 1. The degree of degradation is captured by the parameter *α* with Δ*σ* = *σ*_*b*_(0)*α, E*_*d*_ = *E*_*d*_(0)(1−*α*) and *B*_*d*_ = *B*_*d*_(0)(1 −*α*)^3^. b,c) Contributions of horizontal displacement and of out-of-plane deflection in the strain tensor component *u*_*xx*_ in the degradation region (panel b) and in the external region (panel c). d) Zoom of fold shape for solutions in panel a (from top to bottom *α* = 0, 0.5, 0.6 and 0.7). e-g) Phase diagrams showing fold depth as quantified by *ζ*_*i*_(*l/*2)−*ζ*_*i*_(0) in dependence of the plate bending modulus *B*_*e*_ and degradation degree *α* (panel e), in dependence of pressure *P*_*H*_ and degradation degree *α* (panel f) or in dependence of fold width and degradation degree *α* (panel g). h) Plate profile in the presence of restoring force density *K*_*r*_*ζ*(*x*) (fold degradation *α* = 0.7, *P*_*H*_*/K*_*r*_ = 20 *µ*m). The dashed line shows the corresponding fold profile from panel (a), which does not include restoring forces. e-g) Calculations have been restricted to the regime where strain contributions 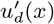 and 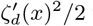 where both below 0.2. Regions that did not fulfill this criterion where shown in white. If not indicated otherwise, parameters were chosen as wing disc length *L* = 200 *µ*m, fold width *𝓁* = 20 *µ*m, basal prestress before degradation start *σ*_*b*_(0) = 0.3 mN/m, *P*_*H*_ = 0.05 kPa, epithelial height *h*_*E*_ = 25 *µ*m, *R* = 15 *µ*m, bending moduli *B*_*e*_ = *B*_*p*_ = *B*_*d*_(0) = 50 × 10^−15^ Nm and 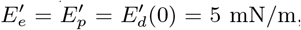, with *E*^*′*^ = *hE/*(1 − *ν*^2^). Apical surfaces and lateral cell boundaries shown in panel a were calculated as described in Appendix C.

Furthermore, we find that softer bending moduli *B*, larger hydrostatic pressures in the cell layer quantified by *P*_*H*_ and bigger fold widths in conjunction with higher degradation degrees *α* promote larger fold depths (*ζ*_*d*_(*l/*2) *ζ*_*d*_(0)), see Fig. 2e-g.

## IV. PLATE DEFORMATIONS IN THE PRESENCE OF A RESTORING ORTHOGONAL FORCE

So far, our solutions to plate shapes were exhibiting out-of-plane deflections which were non-negligible in a larger region of the plate extending significantly beyond the degradation region in the fold, see Fig. 2a. This constitutes a deviation from experimentally observed wing disc fold shapes, see Fig. 4a, Supplemental Movie 3 and^21^. We therefore asked which mechanism might prevent such long-ranged out-of-plane deflections in the biological system. We suggest that one plausible mechanism would be a restoring force that increases with the distance between peripodial membrane and columnar epithelium. Such a mechanism might be realized through spring-like molecular attachments between the apical side of the columnar epithelium and the peripodial epithelium. This attachment might for instance be realized through the proteins of the apical extracellular matrix^32^. In our equations for normal force equilibrium, this restoring force can be incorporated through an additional external force density −*K*_*r*_*ζ*(*x*), where *K*_*r*_ is an effective spring constant. Accordingly, the equation of normal force balance becomes

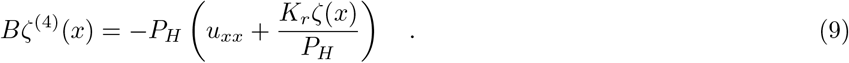

Using this new force balance equation above and assuming again a step-wise increase of Δ*σ* in the degradation region, solutions for the different plate regions can be calculated, see Appendix D. Indeed, we find that plate shape is changed through the restoring force such that out-of-plane deflections are diminished outside the degradation region, while the fold extension in the center remains approximately constant, see Fig. 2h.

*Case of negligible bending rigidity*. In the following, we will consider a special but insightful scenario of negligible bending stiffness in the presence of a smooth degradation profile *α*(*x*) centred in the lower half of the basal plate. (For instance, *α*(*x*) might be a Gaussian curve centered at *x* = 0 in the degradation region with a standard deviation *σ* = *𝓁/*2.) Correspondingly, we may assume Δ*σ*(*x*) = *α*(*x*)*σ*_*b*_(0) and *E*^*′*^(*x*) = *E*^*′*^(0)(1 − *α*(*x*)). In this scenario, orthogonal force balance according to Eq. (9) leads to the simple relation *ζ*(*x*) = −*P*_*H*_*u*_*xx*_*/K*_*r*_. Hence, we obtain that *ζ*(*x*) ≈ −*P*_*H*_ Δ*σ*(*x*)*/*(*E*^*′*^(*x*)*K*_*r*_) and thus

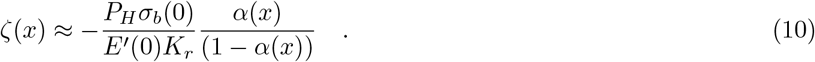

In particular, for small values of *α*(*x*), the fold profile *ζ*(*x*) is proportional to the profile of basement membrane degradation *α*(*x*).

## V. TESTING OF KEY MODEL ASSUMPTIONS IN EXPERIMENTS

Folds observed in the *Drosophila* wing disc presents an ideal model for testing the fundamental assumptions of our theoretical framework. During wing disc development, localized degradation of the basement membrane that is believed to induce the formation of the H/H fold – the fold that demarcates the central boundary of the hinge region^21^.

A key assumption of our model is that fold depth increases with a reduction of basal prestress. This is illustrated by the monotonic dependence of predicted fold depth on the degradation parameter *α* which scales with the decline in basal prestress Δ*σ* = *ασ*_*b*_(0), see e.g. Fig. 2a,d. Vice versa, the model predicts that fold depth decreases in response to an increase in basal prestress. To test this hypothesis experimentally, we manipulated basal prestress in the tip of the wing disc H/H fold by optogenetic recruitment of RhoGEF2, an activator of actomyosin contractility^33^, to the basal surface of the cells in the basal tip of the fold. Optogenetic recruitment was based on the light-dependent binding of Cryptochrome 2 (Cry2) fused to RhoGEF2 (RhoGEF2-Cry2) to a membrane-anchored CIB protein, see^34,35^. As previously described^35^, we illuminated a ‘basal’ volume of cells using two-photon excitation with a pulsed laser of wavelength *λ* = 950 nm for 2 min, see Fig. 3a and Materials and Methods. By this method, the amount of actomyosin-induced active prestress in the basal surface increases. Following the photoactivation, we observed changes in the fold shape and depth. Fig. 3b shows a cross-section of a wing disc before (left panel) and after (right panel) photoactivation. The cells are visualized by Gap43::mCherry (a plasma membrane marker). Fig. 3c illustrates the profile of the basal surface of the folded epithelium, along with labels that illustrate the analysis employed for measuring the fold depth, see Materials and Methods. Notably, right before and after the photoactivation, the fold depth was found to have decreased by ≈4.6%, as illustrated in Fig. 3d,e. This result aligns with our model’s prediction that fold depth decreases in response to an increase in basal prestress.

**Figure 3.**
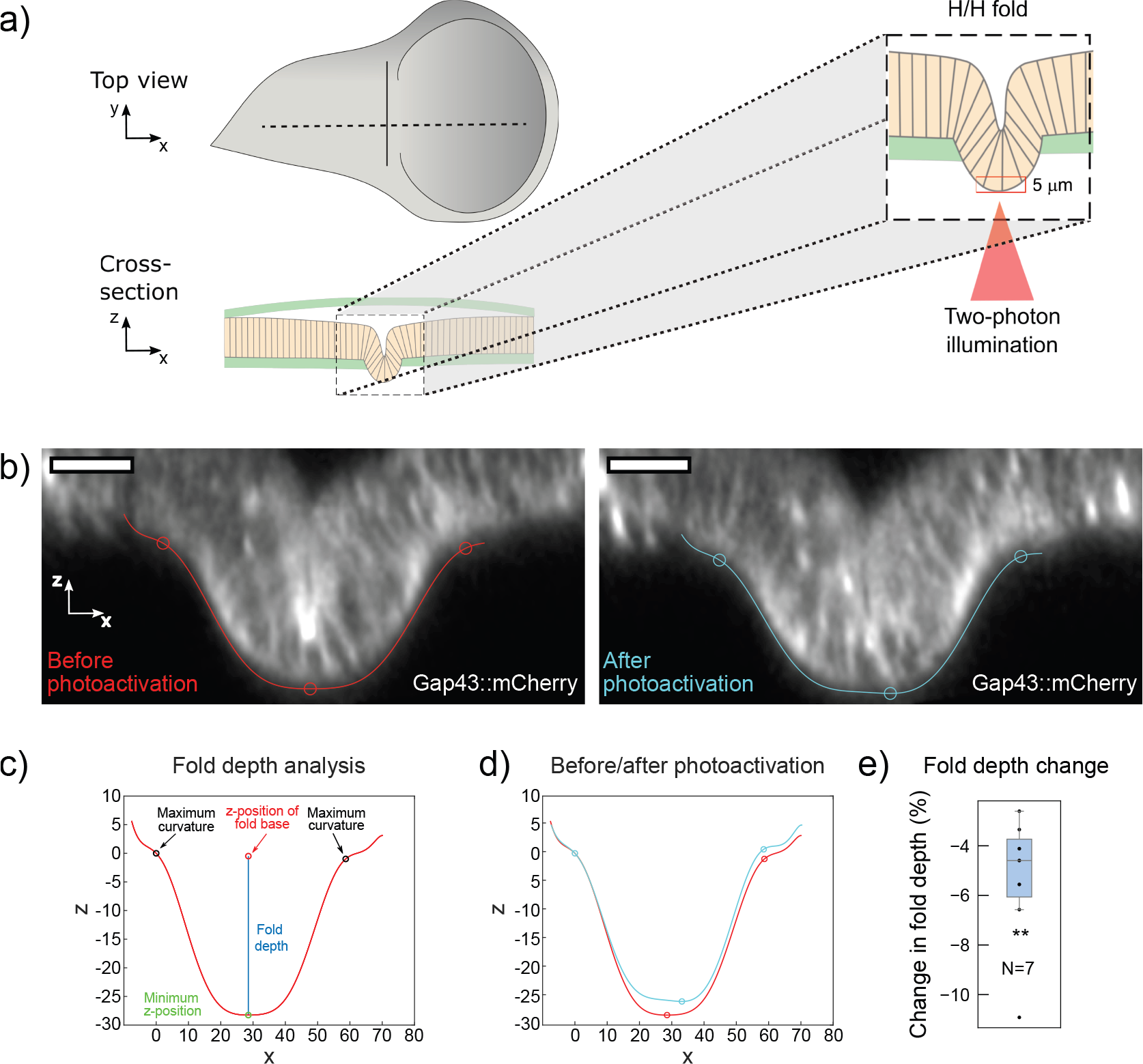
Basement membrane deformation through photoactivation-induced increase of basal tension in third instar *Drosophila* wing disc epithelia of transgenic flies expressing a light-sensitive actomyosin activating construct (RhoGEF2-CRY2), see also Supplemental Movie 2. a) Schematic representation of the top view (upper panel) and the cross-sectional view (bottom panel) of a *Drosophila* wing disc showing the position of the H/H fold. A magnified inset details the cross-sectional view of the epithelial fold with the basement membrane (green) and the location of the targeted area (red box) for photoactivation. b) High-resolution confocal images showing an exemplary single epithelial fold before (left) and right after (right) photoactivation, with cell membranes highlighted by fluorescence (Gap43::mCherry). The red line indicates the initial profile, and the cyan line shows the post-activation profile. Scale bars: 15 *µ*m. c) Fold profile (red) and graphical representation of the fold depth analysis elements, with the initial fold depth marked by the blue line. d) Fold profiles before (red) and after (cyan) photoactivation, indicating the change in the fold depth due to increased contractility at the tip of the fold. e) Box plot detailing the distribution of relative fold depth changes post-photoactivation (N=7). Significance was determined with a one-sample paired T-test against the null hypothesis that the data comes from a normal distribution with a mean equal to zero (∗*p <* 0.05, ∗ ∗ *p <* 0.01, ∗ ∗ ∗*p <* 0.001, n.s. not significant).

**Figure 4.**
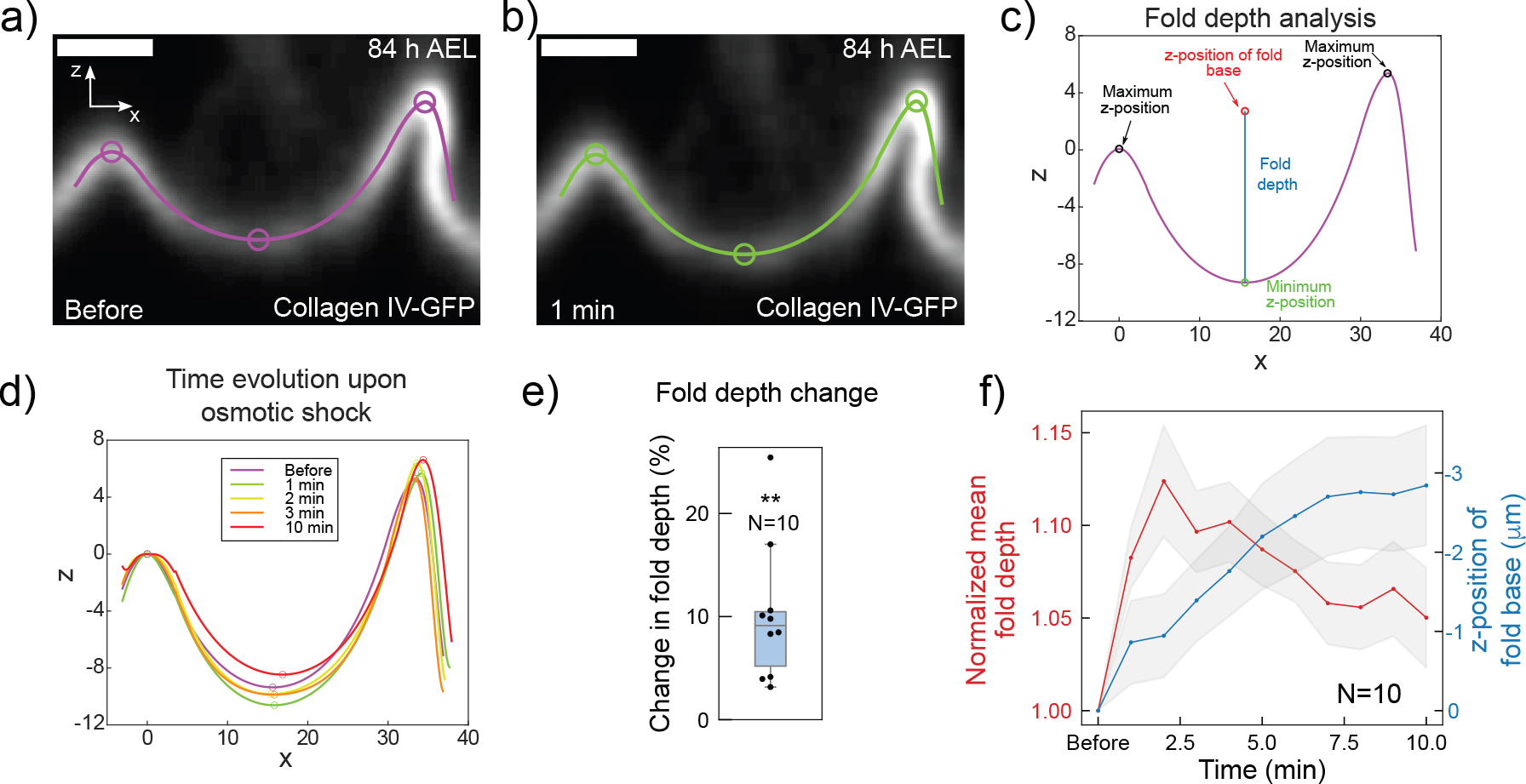
Basement membrane deformation upon hypo-osmotic shock applied to wing disc epithelia of third instar *Drosophila* larvae, see also Supplemental Movie 3. a,b) Time series of confocal microscopy images of the basement membrane (Collagen IV-GFP) of an explanted exemplary wing disc depicting the basal fold profile before (a) and 1 min after water addition to the medium (b). Scale bars, 10 *µ*m. c) Polynomial fit of the cross-sectional profile of a single epithelial fold before the hypo-osmotic shock illustrating the measurement of fold depth (blue line). The end-points of the fold were identified with the points with minimum z-coordinates (black arrows). The fold depth was measured as the distance between the minimum z-coordinate of the fold and the average z-coordinate of the fold end-points. d) Temporal evolution of the fold profile upon hypo-osmotic shock illustrating the fold change over time. e) Box plot representation of relative basal fold depth change of the basement membrane upon hypo-osmotic shock (N=10). Changes were determined by comparing the fold depth before and the average fold depth at three time points right after (*t* = 1, 2, 3 min). f) Mean of normalized fold depth (red curve) and average position of the fold base (blue curve) over time (N=10). Shaded area represents the standard error of the mean for either curve. Time point *t* = 0 min corresponds to measurements before the shock. Wing discs were explanted at developmental stages in the range of 76-96 h AEL. Changes of fold depth were restricted to H/H folds with depth larger than 5 *µ*m. Significance was determined with a one-sample paired T-test against the null hypothesis that the data comes from a normal distribution with a mean equal to zero (∗*p <* 0.05, ∗ ∗ *p <* 0.01, ∗ ∗ ∗*p <* 0.001, n.s. not significant).

One other important aspect of our model is the finding that fold depth increases with hydrostatic pressure excess in the cell, see Fig. 2f. To test that prediction, we performed hypo-osmotic shock experiments on wing discs with small folds of depth smaller than 30 *µ*m dissected from 3rd instar larvae^24^. Shocks were applied by adding 30% of distilled water to the medium, see Materials and Methods. Hypo-osmotic shocks were previously shown to transiently increase the intra-cellular hydrostatic pressure in conjunction with cell swelling^36^. Figs. 4a and b exhibit cross-sectional views of a wing disc expressing Collagen IV-GFP, a major component of the basement membrane, before and immediately after the addition of water to the medium. Fig. 4c illustrates the profile of the basal surface of the folded epithelium, along with labels that illustrate the analysis employed for measuring the fold depth, see Materials and Methods. In accordance with our model, we found that the fold depth increased immediately upon hypo-osmotic shock, see Fig. 4d, with a median increase of ≈ 9%, see Fig. 4e. This fold depth increase was only transiently present during the phase of epithelial swelling, see Fig. 4f. Epithelial swelling is reflected in the increase of mean z-position of the fold base over time.

We conclude that experimental measurements are able to verify key assumptions of our model qualitatively.

## VI. DISCUSSION

Here, we have presented a conceptual study of the mechanics of basement membrane folding upon localized basement membrane degradation in epithelial tissues. We focus on the exemplary case of the larval wing disc tissue of the fruit fly *Drosophila melanogaster* for which folding in conjunction with localized basement membrane degradation has been reported^21^. As the epithelial cytoskeleton is expected to behave fluid-like on developmental time scales, we consider only the solid-like basement membrane as a material scaffold that stores shape memory during epithelial folding. Accordingly, we model the basement membrane as a thin elastic plate. This plate is assumed to be subject to an active cell-generated and a passive elastic prestress. We assume that these two prestresses cancel each other out before local basement membrane degradation starts thereby enabling an initially flat plate conformation in the absence of external forces.

We restrict our study effectively to one space dimension, assuming that the region of degradation is narrow and that the plate profile is constant along the second dimension (y-direction). As localized basement membrane degradation is assumed to be slow as compared to viscoelastic shape relaxation time scales, we solve the problem of plate deformation at increasing basement membrane degradation as a quasi-stationary problem. Furthermore, we regard the wing disc basement membrane plate as a self-balanced continuum of a closed plate structure that comprises a top plate (peripodial) and a bottom plate that are mechanically linked through rigid semi-circular elements at the periphery, see Fig. 1e.

In particular, we find that local plate degradation in a small region of the center of the bottom place can indeed yield local basement membrane folding in the absence of tissue growth. This folding entails a built-in symmetry breaking of folding towards the cell-opposing side which is mediated through either of two mechanisms i) hydrostatic pressure excess in the cells which drives the fold outwards once basal relaxation has started to kick in and ii) top-bottom asymmetry in bending-associated stresses through the presence of the connected basement membrane plate at the top (peripodial).

In our first modelling approach (Section III), orthogonal force balance upon degradation is achieved through contributions of bending stiffness and the external orthogonal force emerging from net hydrostatic pressure as soon as basal cell diameters are widened. Respective solutions of plate deformation show that local plate degradation leads not only to folding in the degradation zone but also to plate deformation at larger distances from this zone. These features make emerging wing disc shape sensitive to the nature of boundary conditions at the wing disc extremities (Fig. 2a). This appears to be a deviation from experimental observations of wing disc fold shapes and compromises robustness and precision of fold positioning in a tissue^21^.

To overcome this model feature, we further introduced a restoring orthogonal force that counteracts the deflection of the basement membrane from its original horizontal position (Section IV). This mechanism could be effectively implemented through a mechanical connection between the apical face of the epithelium and an apical ECM scaffold. We find that this new feature leads to a reduction of basement membrane deflection outside of the degradation zone. In particular, if bending contributions are negligible, we find that the out-of-plane deflection is proportional to the spatial degradation profile for small degrees of degradation, see Eq. (10). Hence, in this parameter regime, epithelial folding at the basal side can be precisely localized, size adjusted and spatially confined through the biological implementation of a localized basement membrane degradation profile.

## VII. MATERIALS AND METHODS

### A Fly stocks and genetics

Flies were raised on standard fly food and maintained at 25°C unless stated otherwise. The following *Drosophila melanogaster* fly stocks were used:

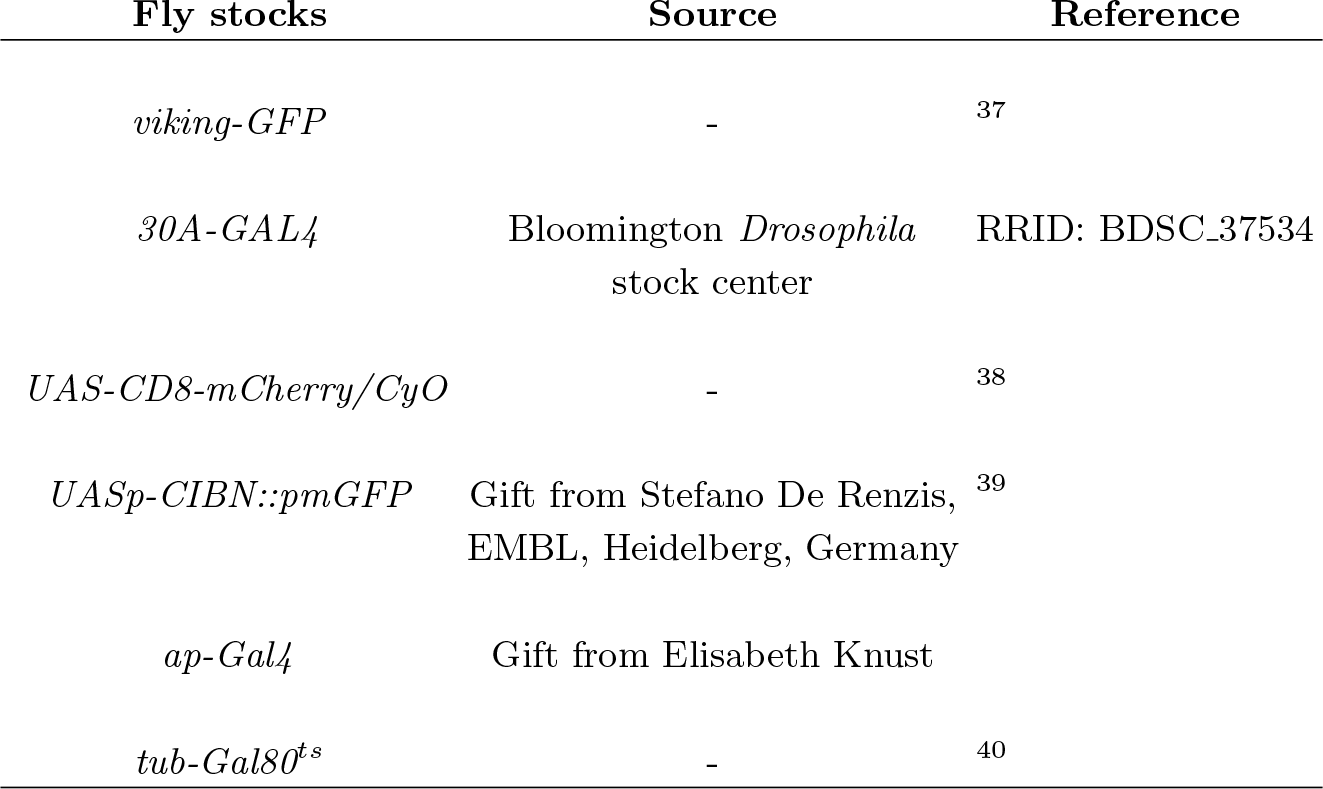

Standard fly husbandry and genetic methodologies were used to cross *Drosophila* strains. The detailed genotype for each experiment were as follows:

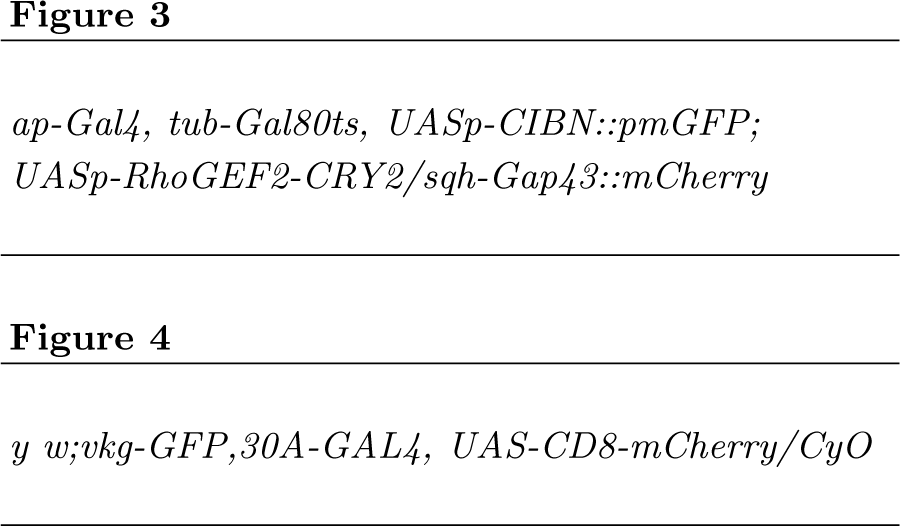

### B. Optogenetically induced tension rise

Incubation, dissection and *in vitro* culture of wing discs from transgenic flies (*ap-Gal4, tub-Gal80ts, UASp-CIBN::pmGFP; UASp-RhoGEF2-CRY2/sqh-Gap43::mCherry*) were done as described previously^21,35^. We note that in this fly line, cell membranes in the wing disc are fluorescently labeled with mCherry. Larvae were incubated in the dark at 25°C and, then, in order to activate expression of the optogenetic construct (UASp-RhoGEF2-CRY2 and UASp-CIBN::pmGFP under the control of ap-Gal4 driver which is inhibited by the temperature-sensitive repressor Gal80ts^34^) transferred to 29°C three days before dissection.

For the experiment, wing discs were dissected in the dark using a red filter^35^ and mounted in a 35 mm glass-bottom petri dish (fluorodish FD35-100) as described before^21^. Then, two-photon activation and imaging were performed via a Zeiss LSM 980 confocal laser scanning microscope, using a C-Apochromat 40x/1.2 water objective.

For every wing disc, the following protocol was applied: initially, wing discs were oriented with the basal side facing the microscope objective. A z-stack of mCherry fluorescence of the wing disc fold region was recorded before photoactivation using a z-interval of 1 *µ*m. Then, photoactivation was performed employing a 950 nm wavelength laser at 8% power for a duration of 2 minutes, targeting the lowermost 5 planes in the stack covering a 5 *µ*m interval at the basal side of the fold. This photoactivation was restricted to a smaller Region of Interest (ROI) in the x-y-plane. This activation ROI was chosen such that the fold was perpendicular to the base of the dish. The x-y extension of the ROI was selected to ensure exclusive photoactivation of the fold’s tip, thereby excluding adjacent areas, see Fig. 3a. Post-photoactivation, a times series of z-stacks of the wing disc fold was recorded in the mCherry channel in the same space extension as before activation comprising in total 5 frames at 2 minute intervals.

For quantitative analysis of the fold depth before and after the two-photon activation, first, the cross section of the wing disc in the fold region of interest is acquired using the reslice command in ImageJ. Then, recorded cross sections are processed by custom MATLAB code that can be downloaded under https://gitlab.com/polffgroup/wingdiscfoldquant (OptogeneticAct FoldDepth analysis.m). In brief, the code first processes the microscopy images by applying a Gaussian blur using ‘imgaussfilt’. Then, the image is thresholded according to fluorescence intensity and binarized. A morphological opening ‘imopen’ is performed to remove smaller objects and holes in the binary image of the fold are filled with ‘imfill’. The edge between the foreground and the background is defined as the profile of the fold. The resulting profile is then smoothed using the function ‘smooth’ and a high order polynomial of degree 10 is fitted over the entire region of interest including the whole fold using the function ‘polyfit’. The curvature of the fitted profile is calculated using a custom MATLAB function LineCurvature2D (https://www.mathworks.com/matlabcentral/fileexchange/32696-2d-line-curvature-and-normals). The points with maximum curvature are taken as the end points of the fold *z*_*max,l*_ and *z*_*max,r*_. Moreover, the minimum z-coordinate *z*_*min*_ of the fitted profile is calculated. The final fold depth is calculated as the difference between the z-coordinates of the fold minimum and the mean of the z-coordinates of the fold edges ((*z*_*max,l*_ + *z*_*max,r*_)*/*2 − *z*_*min*_).

### C. Osmotic shocks

Staging and dissection of wing discs were performed as previously described^21^. Dissection was performed at 76-96 hours After Egg Laying (AEL). The transgenic line used for this experiment expressed fluorescently labeled Collagen IV (vkg-GFP). The cells in the fold region (hinge region) where red-fluorescently labeled by 30A-GAL4, UAS-CD8-mCherry.

Dissected wing discs were carefully placed into cell culture dishes (fluorodish FD35-100) coated with poly-D-lysine (0.1 mg/mL in PBS, Gibco A3890401), see^8^ for details. The wing discs were mounted with their apical side tethered to the glass bottom of the dish.

Imaging, as illustrated in Fig. 4, was carried out using a Zeiss LSM700 confocal microscope at the PoL light microscopy facility. A 40x/1.25 numerical aperture water immersion objective was employed to acquire the images.

First, an initial z-stack of approximately 100 ×100 *µ*m^2^ (1024×1024 pixel^2^) around the fold region was acquired before the hypo-osmotic shock. The size of the stack was chosen to cover the total depth of the fold. The interval between slices was adjusted depending on the developmental stage, with the z-interval ranging from 1 to 1.8 *µ*m. This was done in order to limit light exposure and photo-toxicity.

Subsequently, 600 *µ*L of distilled water (dH_2_O) was added to 2 mL of the culture medium in the dish to reduce the medium’s osmolarity to 77 %. Proper mixing was ensured by using pipette pumping. Following the addition of water, a time series of z-stacks of the region of interest was recorded with 1-minute interval between consecutive images covering ten minutes in total.

For quantitative analysis of fold depths, the microscopy hyperstack is smoothed by applying a Gaussian blur. Then, the cross section of the wing disc is selected and output as a separate time series through the reslice command in ImageJ. In a next step, the depth of the fold was calculated for each time point before and after the hypo-osmotic shock using a custom MATLAB code. (The code can be downloaded under https://gitlab.com/polffgroup/wingdiscfoldquant (OsmoticShock FoldDepth analysis.m).) Within the code, the cross section of the labelled basement membrane is imported and the fold profile is extracted by selecting the pixel coordinates with the maximal intensity of each image column. If the coordinate is not within the fold due to insufficient signal strength, the z-coordinate of the previous point is used. The resulting fold profile is first processed by using a lowpass filter and data smoothing. The end points of the fold are then detected by calculating the two local maxima *z*_*max,l*_ and *z*_*max,r*_ of the profile within the region of interest. The middle part of the fold is fitted with a 4-degree polynomial by using the function ‘polyfit’ and the minimum z-coordinate *z*_*min*_ of the fitted fold profile is calculated. The final fold depth is then calculated as ((*z*_*max,l*_ + *z*_*max,r*_)*/*2 −*z*_*min*_). The absolute value of the average of the z-coordinates *z*_*max,l*_ and *z*_*max,r*_ of both end points also accounts for the fold base, see Fig. 4f.

We note that the analysis of osmotic-shock-induced changes of wing disc folds slightly differs from our analysis of changes of wing disc folds during optogenetic experiments. This was required since the fly lines in the two experiments were different in fluorescent cell-labeling and wing disc morphology.

### D. Supplementary Movie 1

Basement membrane plate deformation over time corresponding to data presented in Fig. 2a.

### E. Supplementary Movie 2

Cross-sectional view of a cultured wing disc from *Drosophila melanogaster*, captured before and immediately after a 2 minute two-photon excitation. The video showcases fluorescence intensity of Gap43::mCherry which is a membrane marker. An initial acquisition is performed before activation (frame 1), followed by a 2 minute excitation period (indicated by a white second frame in the movie), and then continuous acquisitions were recorded at 2 minute intervals after the activation. Scale bars represent 15 *µm*.

### F. Supplementary Movie 3

Cross-sectional view of an 84 h AEL (After Egg Laying) cultured wing disc of *Drosophila melanogaster* before and 1 minute after the addition of water to the medium, inducing a mild hypo-osmotic shock. The dynamics of the fold shape change were captured by taking z-stacks in 1 min intervals. The video visualizes Collagen IV-GFP (green) marking the basement membrane and 30A-GAL4, UAS-CD8-mCherry (magenta) highlighting hinge cells. Scale bars represent 10 *µm*. Note that the tissue deswells after ≈15 min due to active cell volume regulation.

## ACKNOWLEDGMENTS

CD and EFF acknowledge financial support from the Deutsche Forschungsgemeinschaft under Germany’s Excellence Strategy, EXC-2068-390729961, Cluster of Excellence Physics of Life of TU Dresden. Furthermore, EFF was supported by the Deutsche Forschungsgemeinschaft (DFG, German Research Foundation) by the grant FI 2260/9-1 and by the Heisenberg program – project number 495224622 (FI 2260/8-1). In addition, the authors thank the CMCB Light Microscopy Facility and the PoL Light Microscopy Facility for excellent support.

## Appendix A: Force and torque balance for thin curved surfaces

Let Γ be a two-dimensional curved surface with parametrisation **X** : ℝ^**2**^ → **Γ** and corresponding tangent space basis vectors **e**_**1**_, **e**_**2**_ and surface normal **n** = **e**_**1**_ × **e**_**2**_*/*|**e**_**1**_ × **e**_**2**_|. We denote forces and torques in the plate by 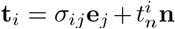 and 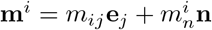 respectively^41^. Furthermore, external forces are given by 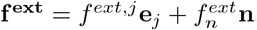 Tensions and torques in the plate arise from bending contributions, surface tension, as well as elastic contributions. In steady state, the following defining force balance equations must hold^41–43^

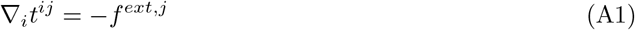

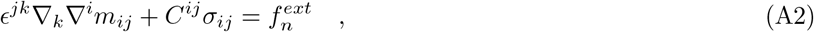

where *C*_*ij*_ = −**n** · *∂*_*i*_**e**_*j*_ is the curvature tensor. The tension contribution from elastic in-plane deformation is^26^

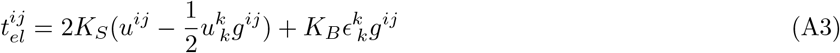

where *K*_*S*_ and *K*_*B*_ are the area shear modulus and the area bulk modulus of the plate, respectively, and *u*_*ij*_ are the components of the elastic strain tensor. Tension contributions from plate bending can be neglected to first order due to a flat reference configuration^43^.

In the unperturbed state, we anticipate that our plate’s extension is in the x-y plane (Fig. 1c). For the special case that i) external tangential net forces are zero, ii) stress, strain and deformation components are constant along the y-axis apart from a possible linear change of *u*_*y*_, and iii) *C*^*xy*^ = *C*^*yy*^ = 0, we obtain as defining equations^25^

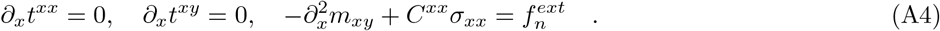

We conclude that *σ*_*xx*_ and *σ*_*xy*_ are constant. As *σ*_*xy*_ ∝*u*_*xy*_ = *∂*_*x*_*u*_*y*_*/*2, we may infer that *u*_*y*_ = 2*u*_*xy*_*x* + *c*_1_(*y*). Correspondingly, *u*_*yy*_ = *∂*_*y*_*c*_1_(*y*), i.e. *u*_*yy*_ must be constant as we proclaimed no y-dependence of stress and strain components.

Let the function *ζ*(*x*) denote the displacement of the neutral line of the plate in z-direction. Then the curvature of the plate *C*^*xx*^ is to first order given by − *ζ*^*′′*^(*x*). Correspondingly, the bending moment *m*_*xy*_ is *m*_*xy*_ = − *Bζ*^*′′*^(*x*), where *B* is the plate’s bending modulus. We therefore obtain

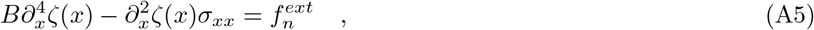

which also follows from the Föppl-von Kármán equations for the equilibrium of plates with large deflection^25,27^.

## Appendix B Boundary conditions

## Appendix C: Estimating the shape of the apical surface

In the following, we describe how the out-of plane deflection 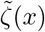 of the apical surface is estimated as shown in Fig. 2a. The local change in cell volume is given by

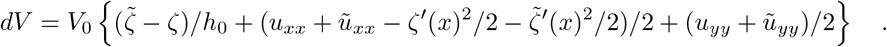

We anticipate, that the apical extension of cell surfaces along the wing disc long axis does not change in the folding process as was reported previously from experimental observations^21^, i.e. *ũ*_*xx*_ ≈0. We further assume that *ũ*_*yy*_ = *u*_*yy*_ to keep equal y-coordinates of basal and apical cell edges.

To find an estimate of the apical cell surface shape in the degradation region, we choose to calculate 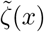 by minimizing cellular volume changes in the degradation region, i.e. minimizing the functional

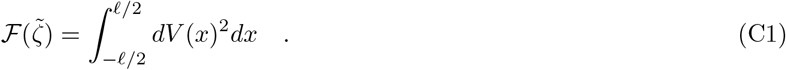

To obtain an approximate numerical solutions for the minimizing function 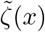, we make the ansatz of a polynomial expansion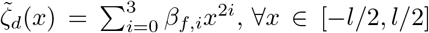, ∀*x* ∈ [−*l/*2, *l/*2]. Here, coefficients *β*_*f,i*_, *i* = 1, …, 3 are determined through the minimization of 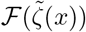 under the constraint that 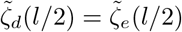 and 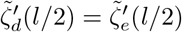 Outside of the degradation *d e* region, we approximate 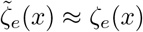

Lateral cell boundaries are drawn as straight lines connecting the points (*x*_*i*_ + *u*_*d*_(*x*_*i*_), *ζ*_*d*_(*x*_*i*_)) and 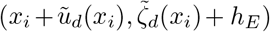, with *x*_*i*_ = *i*Δ*x, i* = 0, ±1,± 2, … and Δ*x* = 2.5 *µ*m. This value of Δ*x* was motivated by typical cell diameters in the wing disc in the young 3rd instar larva.

## Appendix D: Solution in the presence of a restoring force

The governing equation in the situation where y-deformations are negligible are

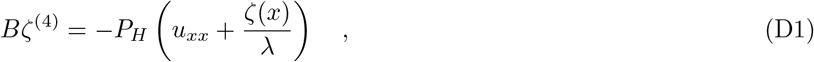

with *λ* = *P*_*H*_*/K*_*r*_. Assuming that Δ*σ* is non-vanishing only in the degradation region, *u*_*xx*_ = Δ*σ/E*_*d*_*′* and zero outside, we first find solutions *ζ*_*h*_(*x*) that solve the homogeneous differential equation

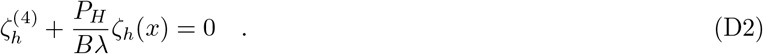

A full solution is then given by *ζ*(*x*) = *ζ*_*h*_(*x*) − *λu*_*xx*_. We make the ansatz *ζ*_*h*_(*x*) = exp(*krx*), where 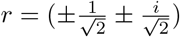 and 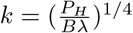

We define the following antisymmetric real-valued functions

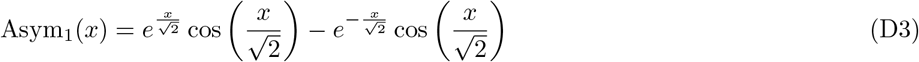

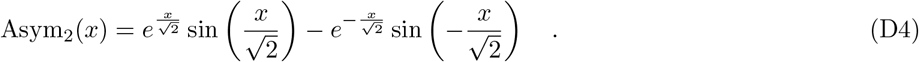

Further, we define the following symmetric real-valued functions

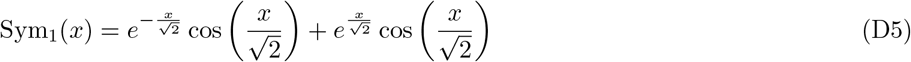

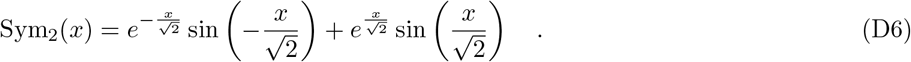

Accordingly, we make the following ansatz for the solution of out-of-plane deflections in the degradation region, the external region in the columnar epithelium and in the peripodial membrane

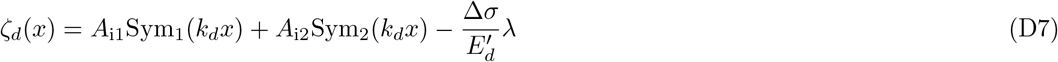

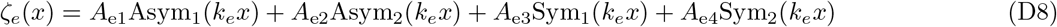

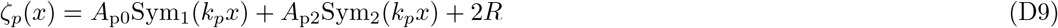

In-plane displacements *u*(*x*) follow from out-of-plane deflections via Eqn. (8). Constants of integration are fixed by boundary conditions 1-9 in Table I.

